# Structural analysis of the Spike of the Omicron SARS-COV-2 variant by cryo-EM and implications for immune evasion

**DOI:** 10.1101/2021.12.27.474250

**Authors:** Dongchun Ni, Kelvin Lau, Priscilla Turelli, Charlene Raclot, Bertrand Beckert, Sergey Nazarov, Florence Pojer, Alexander Myasnikov, Henning Stahlberg, Didier Trono

## Abstract

The Omicron (B.1.1.529) SARS-COV-2 was reported on November 24, 2021 and declared a variant of concern a couple of days later.^1,2^ With its constellation of mutations acquired by this variant on its Spike glycoprotein and the speed at which this new variant has replaced the previously dominant variant Delta in South Africa and the United Kingdom, it is crucial to have atomic structural insights to reveal the mechanism of its rapid proliferation. Here we present a high-resolution cryo-EM structure of the Spike protein of the Omicron variant.

## RESULTS

The Omicron Spike cDNA was obtained by reverse transcription of a nasopharyngeal viral sample and cloned into an expression vector as previously described.^3^ The protein was purified to homogeneity *via* its TwinStrep tag from the supernatant of transiently transfected ExpiCHO cells (Fig. S1), and vitrified for cryo-electron microscopy (cryo-EM) at a concentration of 0.35 mg/ml as described in the Methods. Grids were screened with a 200kV TFS Glacios cryo-EM, and final data collection was performed on a 300kV TFS Titan Krios G4. Data was processed ‘on-the-fly’ with CryoSPARC Live, resulting in a first 3D map after 3 hours from the Glacios.^4^ A total of 8672 dose-fractionated electron event recordings (movies) were then collected on the Titan Krios G4, processed first with cryoSPARC Live and re-processed manually with cryoSPARC version 3.3.1. The resulting map of the full Omicron Spike was resolved at a resolution of 3.02 Å (FSC 0.143 criterion) with C1 symmetry (Fig. S2). A map with C3 symmetry was determined to a resolution of 2.65 Å. A local mask on a region containing both a receptor binding domain (RBD) and a N-terminal domain (NTD) was applied. Focused refinement specifically on this masked region yielded a map at 3.88 Å resolution.

The cryo-EM maps of the Omicron Spike glycoprotein clearly revealed many of the mutations present in this variant (Fig. S3). The overall architecture of the Spike trimer was comparable to the first structure of the wild-type Spike (PDB ID 6VSB) (RMSD CD all atoms 2.4 Å) with one RBD in the up position and two RBDs down (Fig. 1a-b)^5^, although a minority of Spike trimers were in a closed conformation with all RBDs down. We did not observe other populations containing two or three RBDs in the up configuration. This suggests that the Omicron Spike may preferentially be in either a one RBD-up or in a closed state (Fig. S2). The best-defined mutations are within the S2 domain intercalating between Spike monomers and within the NTD (Fig. 1c, e). A view from the top of the Spike protein exposes a large cluster of mutations on the RBDs (Fig. 1d-e).

**Figure 1:**
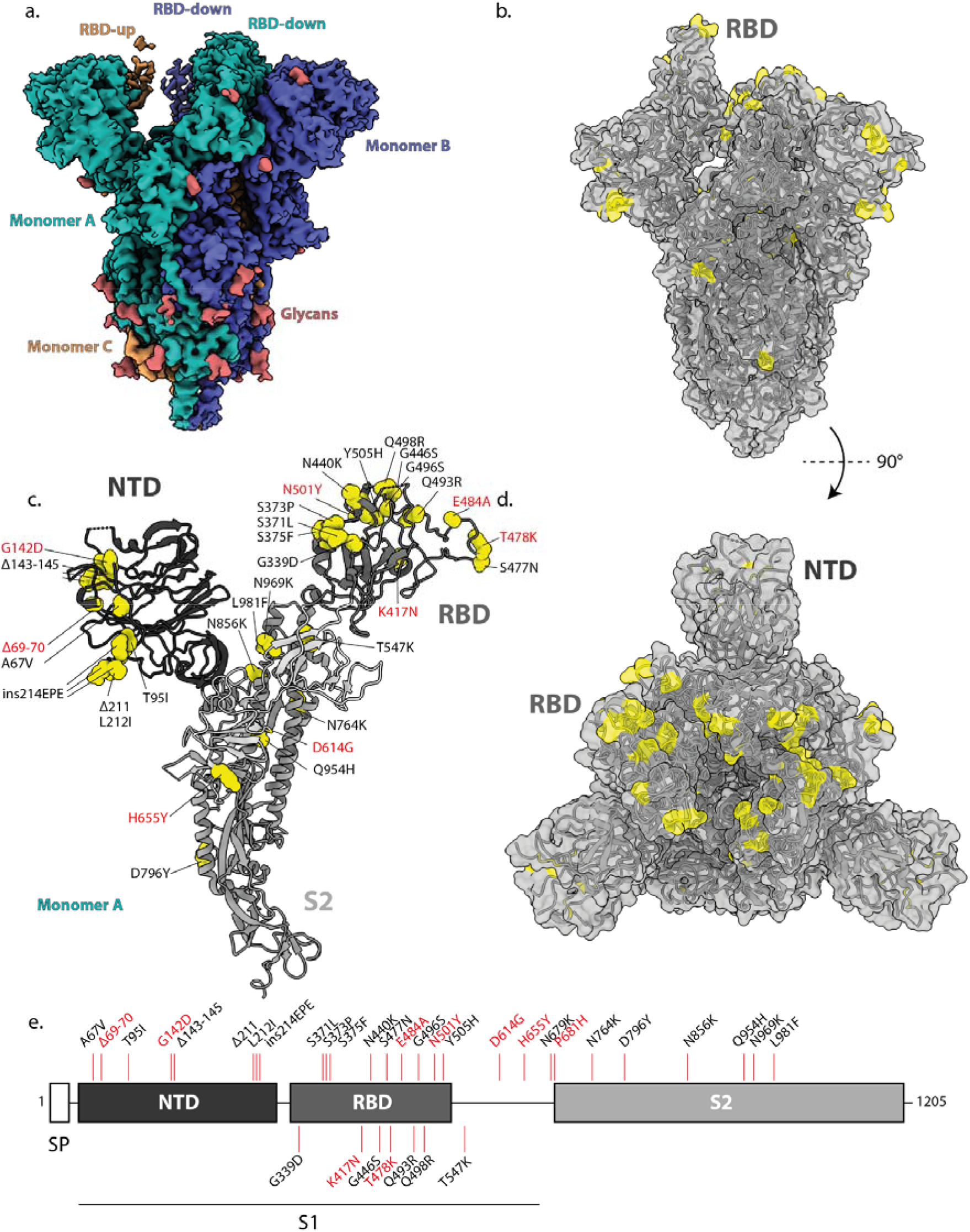
Cryo-EM map and model of Spike Omicron Variant. a. Cryo-EM map of the Spike of the Omicron variant. Map coloring corresponds to each of the Spike monomer chains that comprise the full trimer (A (green), B (blue) and C (orange)). Red indicates a glycan. The flexibility of the single RBD-up (monomer C) is barely visible. b. Side-view surface representation of the atomic model of the Omicron Spike in gray, with mutations highlighted in yellow. c. Ribbon representation of monomer A, highlighting the different domains colored in gray-scale (as in panel d) and the mutations in spheres in yellow. Mutations are labelled. Red labelled mutations are those shared with other VOCs. d. Top view of panel b of the Omicron Spike, with specific mutations highlighted in yellow. e. Overview of the domain architecture of the Omicron Spike. Specific domains are highlighted: signal peptide (SP), N-terminal domain (NTD), receptor binding domain (RBD), S1 and S2 domain.

The exceptional quality of the map in the S2 domain reveals potential new interactions formed by Omicron-unique mutations. The D796Y (all numbering based on wild-type positions) mutation replaces a charged surface-exposed acidic residue with tyrosine, an amino acid that contains an aromatic sidechain (Fig. 2a). We observe that Y796 allows for potential carbohydrate-pi interactions with the N-linked glycan chain originating from N709 of the neighboring monomer chain.^6,7^ We suspect that this interaction could have a stabilizing effect for the spike trimer.

**Figure 2:**
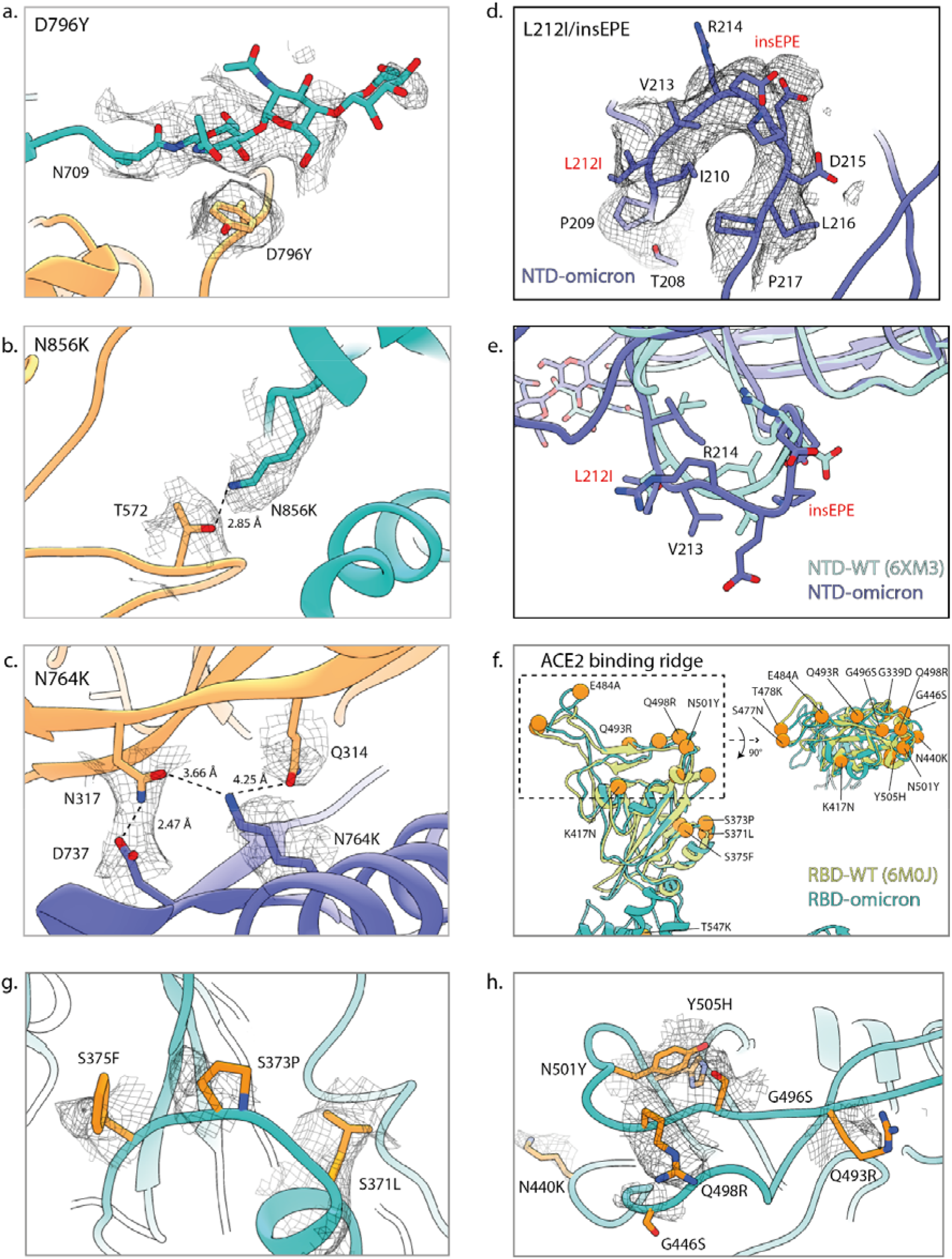
Omicron mutations alter local conformations and forms new interactions. Zoomed in ribbon representations of the Omicron spike at sites of mutations. Mesh represents the cryo-EM map. Colors of models represent the Omicron Spike monomer chain as colored in Fig. 1a. a. D796Y mutation allows for the potential carbohydrate-pi interactions with N-linked glycans from N709 of a neighboring Spike monomer. b. N856K mutation of the fusion peptide. The longer side-chain of the lysine residue allows for a new interaction with T572 from an adjacent monomer. c. The N764K mutation forms a new network of interactions with both N317 and Q314 from an adjacent monomer. d. NTD mutation at a loop that contains a deletion at residue 211, a mutation L212I and an insertion of EPE at position 214. Numbering is based on that of the original wildtype Spike. e. Superposition of the same loop depicted in panel d. NTD-Omicron (dark blue) to the wild-type NTD (light blue) shows a large change in register and chemical environment of the loop between amino acids 210-215, wild-type numbering. f. Superposition of the RBD-Omicron (green) to the wild-type RBD (light green). Highlighted in the dashed-box is the receptor binding region that interacts with ACE2. Mutations present on the Omicron RBD are highlighted as orange spheres. g. Omicron RBD mutations represented at sticks (orange) centered on the mutations S371L, S373P, S375F. h. Omicron RBD mutations represented at sticks (orange) centered on the N501Y mutation.

The N856K mutation is within the fusion peptide of the Spike protein.^8^ The fusion peptide is required for attachment and anchoring of the virus directly to a host’s cell membrane. It is exposed during proteolytic cleavage that liberates the S1 from the S2 domain followed by subsequent cleavage at the S2’ site. We observe the N856K mutation with the longer side-chain of the lysine residue able to form new interactions with T572 from an adjacent monomer (Fig. 2b). We can only speculate that this may have a functional impact on the release of the fusion peptide. In addition, the mutation from an asparagine to lysine may aid cell membrane penetration.^8–10^

Other mutations located within S2 with stabilizing potential, such as N764K, can also be clearly visualized (Fig. 2c). We can model an amine head-group of K764 forming two new interactions with Q314 and N317 from an adjacent monomer, which could stabilize the Spike trimer. In particular, Q314 and N317 precede the RBD in sequence (residues 330-520) and are just below the linker that allows for RBD transition from a down to an up position.

We observe that the RBD is predominantly in a 1-up or closed state, allowing for the Spike to occlude efficiently RBD epitopes. Thus, the many mutations present in the S2 domain present only in the Omicron Spike protein may allow for optimization of RBD positions that allow for receptor binding while concealing epitopes that are the targets of neutralizing antibodies.^11^

The NTD is the target of various neutralizing antibodies and constitutes an antigenic supersite,^12–14^ and was subjected to focused refinement (Fig. 2d-h). We observe that the overall fold of the Omicron NTD is relatively unchanged compared to either its wild-type (RMSD 1.7 Å) or Delta (RMSD 2.4 Å) counterparts (Fig. S4a-b). The loop encompassing deletions 143-145 (N3 loop)^12–14^ has a different conformation compared to the Delta NTD. The Delta variant Spike also contains NTD mutations in the same N3 loop (E156G, Δ157-158). Superpositions of structures of Fab-bound NTDs predict that the Omicron NTD is resistant to some NTD antibodies due to steric clashes and changes in the antibody binding surface (Fig. S4c).^12–14^ We identify a second loop (residues 208-217), hereby called the EPE loop, which contains Omicron-specific mutations, deletions and inserts: deletion 211, L212I and insertion at 214 of EPE (Fig2. d-e). The locally-refined cryo-EM map in this region allowed for the *de novo* building of this loop with the aid of a model generated from AF2.^15^ In the Omicron NTD, the EPE loop follows the C□ backbone as the wild-type NTD, however the chemical nature is drastically changed with the introduction of two acidic negatively charged residues (Fig. 2e). This insertion is not predicted to affect the binding of known NTD targeting antibodies, being localized far from the site of antibody binding (Fig. S4c).^12,13^

The RBD is the target for a vast number of neutralizing antibodies elicited by vaccination and previous infection. All commercially available antibodies that have been approved for clinical use, target this domain of the Spike protein. Based upon previous variants of concern (VOCs) and experimental studies, some of the mutations such as N501Y and Q498R may independently or synergistically enhance binding of the Spike to its receptor, the human ACE2 protein.^16,17^ In addition, the mutations that decorate the RBD may allow for immune evasion or render monoclonal antibody therapy ineffective.^18–20^ Alignment of the wild-type RBD with that of the Omicron reveals only slight changes with a RMSD of 2.6 Å (Fig. 2f). The locally-refined map in this region allowed us to model all the mutations located within the RBD (Fig. 2g-h and S3). A region that encompasses residues 371 to 375 contains three serine residues mutated: S371L, S373P, S375F. These residues are on a surface exposed region of the RBD and may play a role in immune evasion of the Omicron variant. In addition, the top saddle of the RBD (Fig. 2f) is the location of many mutations shared with others VOCs, in particular N501Y that has been shown to foster ACE2 binding, along with other Omicron-specific mutations such as Q493R and Q498R.^16,17^ Overall the Omicron RBD has a highly mutated surface that encompasses the epitopes of the known major classes of antibodies (Fig. S5a). Superpositions of structures of representative Fab-bound RBDs suggest that all 4 classes of RBD targeting antibodies have their ability to bind abrogated (Fig. S5b).^21–27^ These observations suggest that the Omicron variant has evolved immune evading mutations from various classes of RBD-targeting antibodies.

Many of the RBD mutations are also located along a ridge that interacts directly with the helices of the ACE2 receptor (Fig. 2f). We performed biolayer interferometry (BLI) binding assays to evaluate the affinity of ExpiCHO-expressed dimeric Fc-human ACE2 to full-length Omicron Spike (Fig. S6). Compared to the wild-type Spike, hACE2 binds to the Omicron Spike with similar affinities and comparable to that of binding to the Delta Spike (4.4 nM vs. 6.8 nM vs. 6.1 nM; wild-type vs. Omicron vs. Delta). The Omicron Spike also shares many RBD-specific mutations (namely N501Y, K417N and mutation at E484) with other VOCs, namely Alpha, Beta and Gamma. These variants all show relatively stronger binding to hACE2 than Omicron (Fig. S6). We conclude that the appearance of a large number of mutations at the ACE2 binding interface do not significantly alter the overall binding affinity of dimeric Fc-hACE2 to the Omicron Spike. In addition, we tested binding to the dimeric mouse ACE2 receptor as it has been previously reported that N501Y containing VOCs had increased ability to infect mice and bind ACE2.^28–30^ The dimeric Fc-mACE2 displays robust binding to the Omicron Spike with an almost 8-fold increase in binding affinity compared to wild-type. Mouse ACE2 also binds significantly stronger compared to other variants with the exception of gamma.

## DISCUSSION

Here we present a high-resolution structure of the Spike protein of Omicron, which is predicted to soon become the dominant SARS-CoV-2 variant worldwide (as of December 2021). The overall structure of the full-length Spike, its RBD and NTD domains are similar to that of the wild-type but with specific changes in both local conformations and surfaces involved in antibody recognition. The location of the mutations in both the RBD and NTD suggests that there has been a strong evolutionary pressure on the Omicron Spike to evade the humoral immune response. The high-quality maps in parts of the S2 domain allowed us to visualize Omicron-specific mutations that form new stabilizing interactions within the Spike trimer, and which may optimize its ability to engage and bind ACE2, while occluding epitopes that are targets of neutralizing antibodies.^11^ Even with a heavily mutated ACE2-binding interface, our binding experiments show that the Omicron RBD is capable of interacting with human ACE2 receptor as efficiently as that of the Delta variant. In addition, the robust binding of mouse ACE2 to the Omicron Spike suggests an increased ability to infect other animal species such as rodents.

Our structure highlights the many mutations found on the Omicron Spike, which we speculate may affect protein stability, glycan conformation diversity, membrane fusion, ACE2 and importantly evasion from antibodies from vaccination or previous infection.

During the completion of this manuscript a complementary preprint describing the Omicron Spike was released.^31^

**Table 1:** List of Mutations of VOC

| **Variant of Concern** | **Lineage** | **List of Mutations** |
| --- | --- | --- |
| **Omicron - o** | B.1.1.529 | A67V, Δ69-70, T95I, G142D, Δ143-145, Δ211, L212I, ins214EPE, G339D, S371L, S373P, S375F, K417N, N440K, G446S, S477N, T478K, E484A, Q493R, G496S, Q498R, N501Y, Y505H, T547K, D614G, H655Y, N679K, P681H, N764K, D796Y, N856K, Q954H, N969K, L981F |
| **Alpha - α** | B.1.1.7 | Δ69-70, Δ144, N501Y, A570D, D614G, P681H, T716I, S982A, D1118H |
| **Beta - β** | B.1.351 | L18F, D80A, D215G, Δ242-244, R246I, K417N, E484K, N501Y, D614G, A701V |
| **Gamma - γ** | P.1 | L18F, T20N, P26S, D138Y, R190S, K417T, E484K, N501Y, D614G, H655Y, T1027I, V1176F |
| **Delta - δ** | B.1.617.2 | T19R, G142D, E156G, Δ157-158, L452R, T478K, D614G, P681R, D950N |

**Table 2:** Cryo-EM collection and data processing

|  | SARS-CoV-2 S Omicron Spike B.1.1.529<br>Full map | SARS-CoV-2 S Omicron Spike B.1.1.529<br>Local map (NTD and RBD) |
| --- | --- | --- |
| **Microscope** | TFS Titan Krios G4 |  |
| Detector | Falcon IV |  |
| Magnification | 96kx |  |
| Voltage (kV) | 300 |  |
| Electron exposure (e-/Å<sup>2</sup>) | 60 |  |
| Defocus range (μm) | -0.8 to -1.8 |  |
| Pixel size (Å) | 0.83 |  |
| Symmetry | C1 | C1 |
| Micrographs | 8,672 | 8,672 |
| Initial particle images (No.) | 470,735 |  |
| Final particle images (No.) | 76,450 | 76,450 |
| Map resolution (Å) | 3.02 | 3.88 |
| FSC threshold | 0.143 | 0.143 |
| **Refinement** |  |  |
| Initial model used | 6MXL | --- |
| Map sharpening B factor (Å<sup>2</sup>) | 42 | 54 |
| Protein residues | 3,295 | 1,166 |
| RMSD deviations |  |  |
| Bond lengths (Å) | 0.002 | 0.003 |
| bond angles (°) | 0.518 | 0.631 |
| **Validation** |  |  |
| MolProbity score | 1.77 | 2.06 |
| Clashscore | 6.43 | 9.73 |
| Poor rotamers (%) | 0.07 | 0 |
| **Ramachandran plot** |  |  |
| Favored (%) | 93.81 | 90.04 |
| Allowed (%) | 5.98 | 9.7 |
| Disallowed (%) | 0.21 | 0.27 |
| **PDB** | **7QO7** | **7QO9** |
| **EMDB** | **14086** | **14087** |

## SUPPLEMENTARY FIGURES

**Figure S1:**
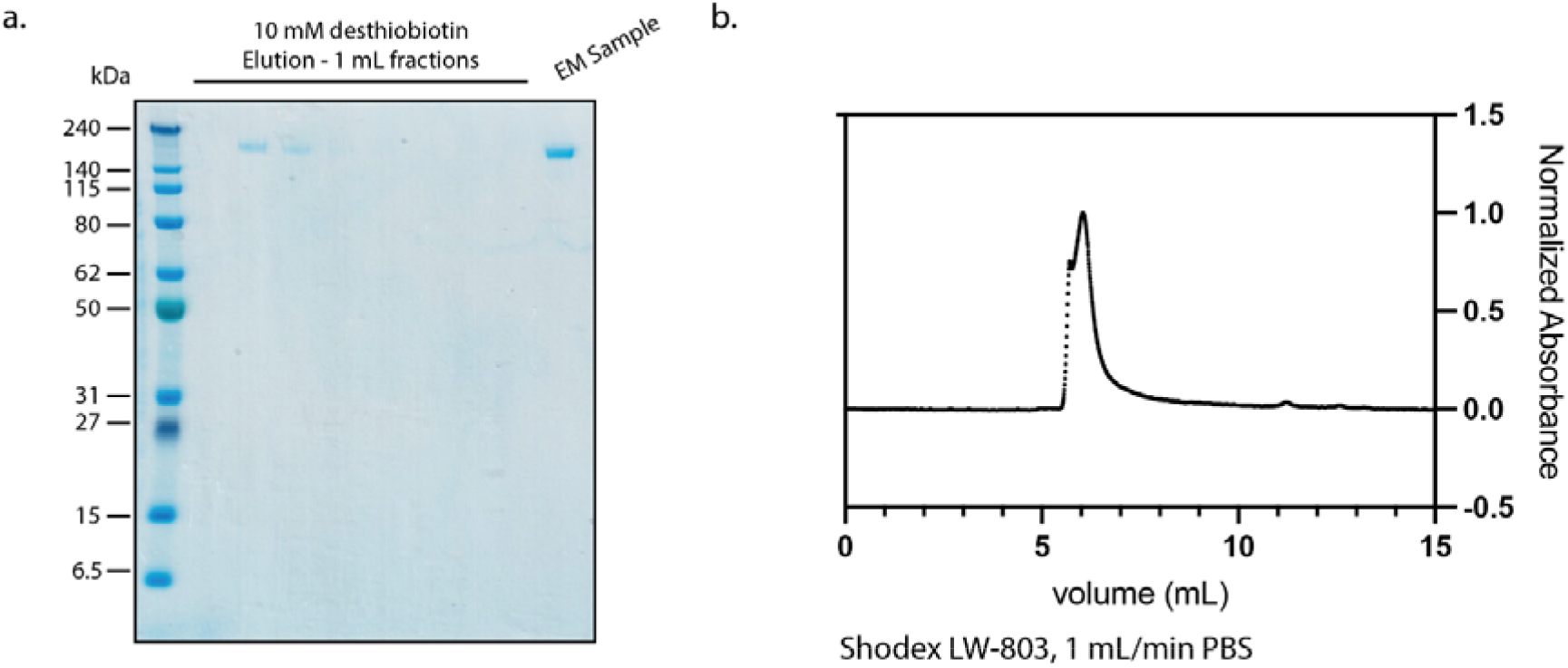
Purification of Spike Omicron variant. a. SDS-PAGE gel of full-length Spike Omicron purified from ExpiCHO cells; purification by StrepTrap HP and final sample used for cryo-EM studies b. Size exclusion chromatography of Spike Omicron.

**Figure S2:**
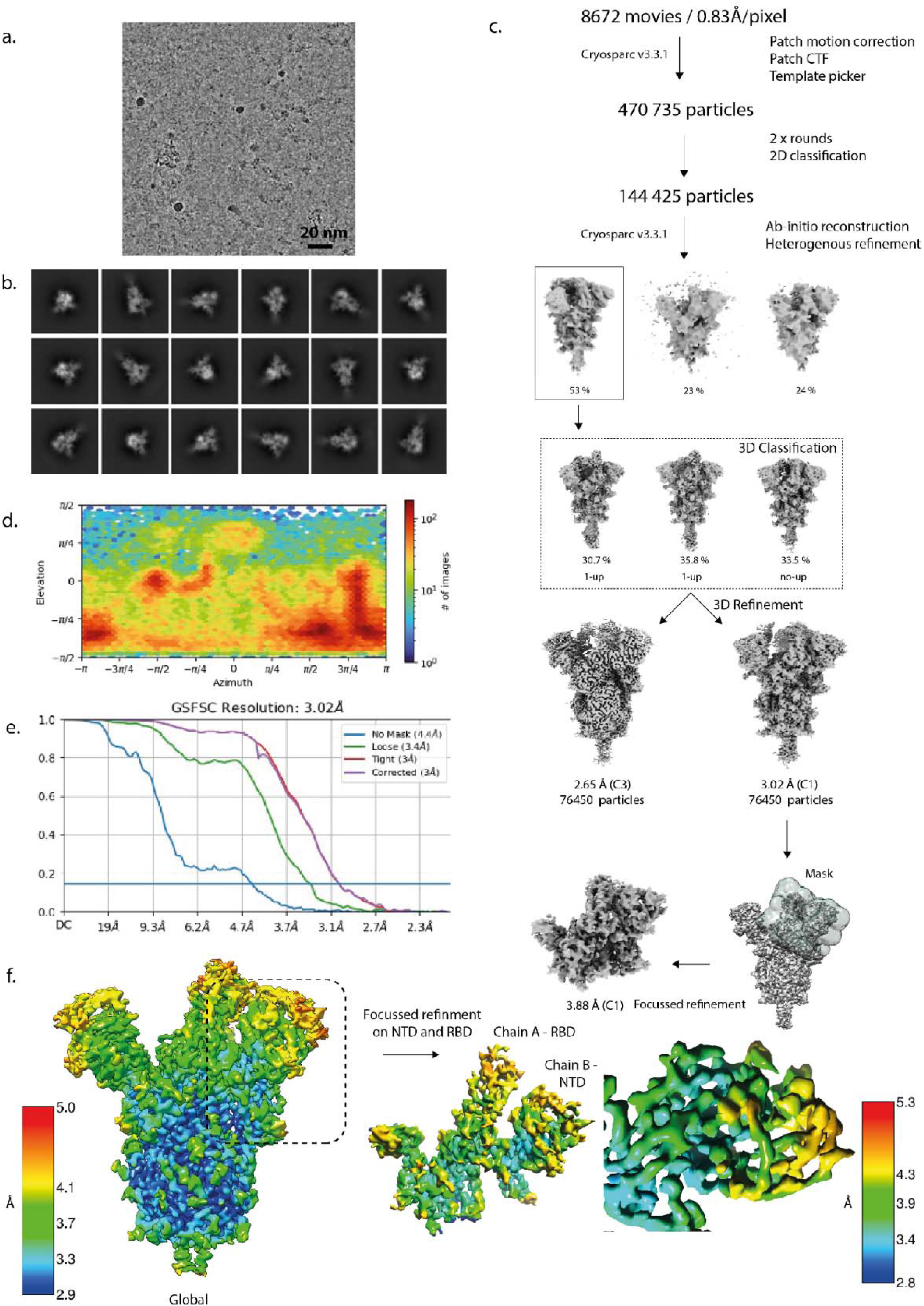
Cryo-EM processing of the Omicron Spike. a. Raw representative micrograph. b. Representative 2D class averages. c. Cryo-EM processing workflow performed in CryoSPARC. d. Direction distribution plot. e. FSC curves indicating a resolution of 3.02 Å (FSC 0.143). f. Final global and focused refined maps colored by local resolution.

**Figure S3:**
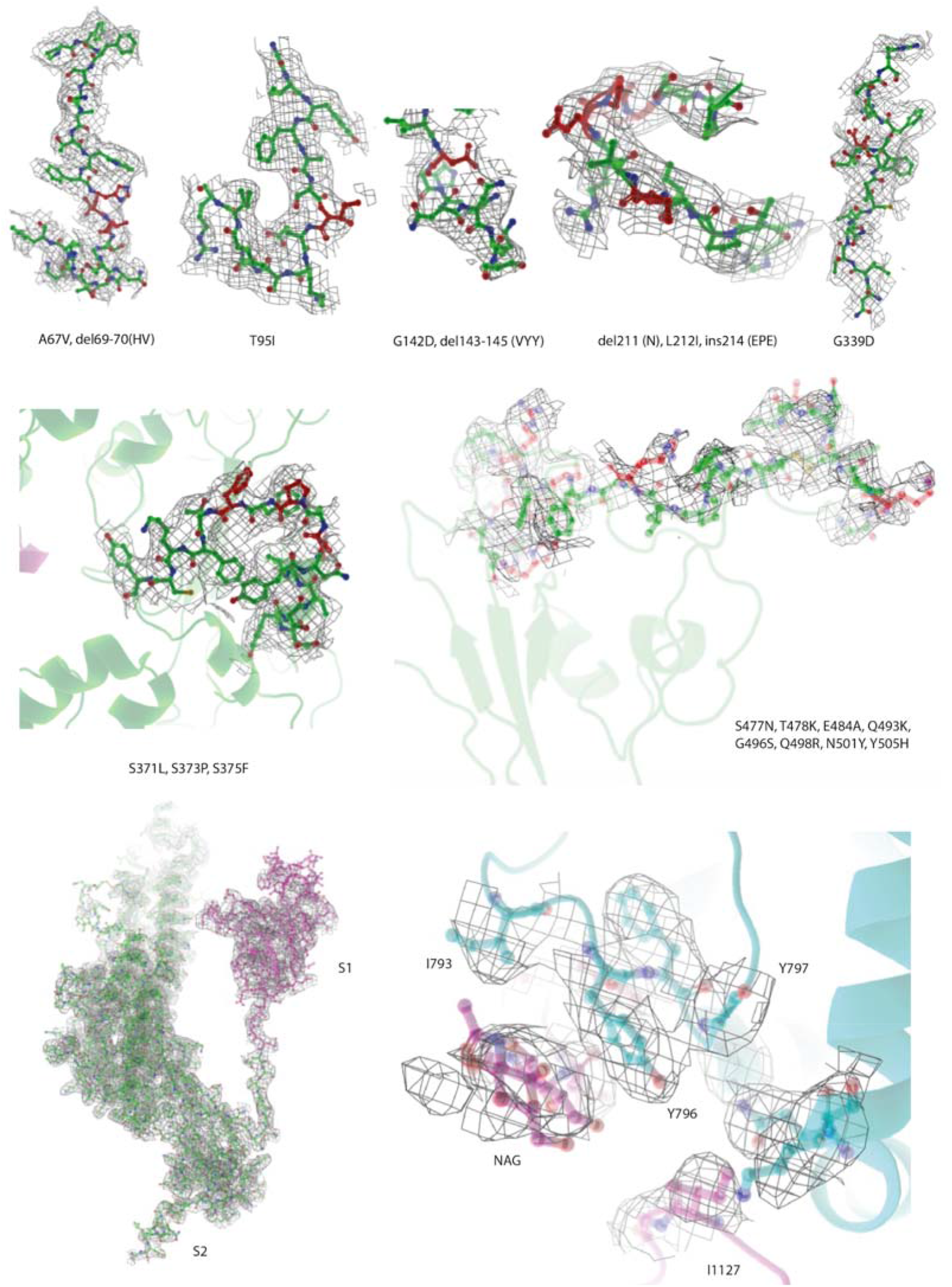
Highlights of regions containing mutations of the Omicron Spike with cryo-EM density maps. The atomic model is shown as ribbon or atom representation. Mutations present in the Omicron Spike are indicated and colored in red.

**Figure S4:**
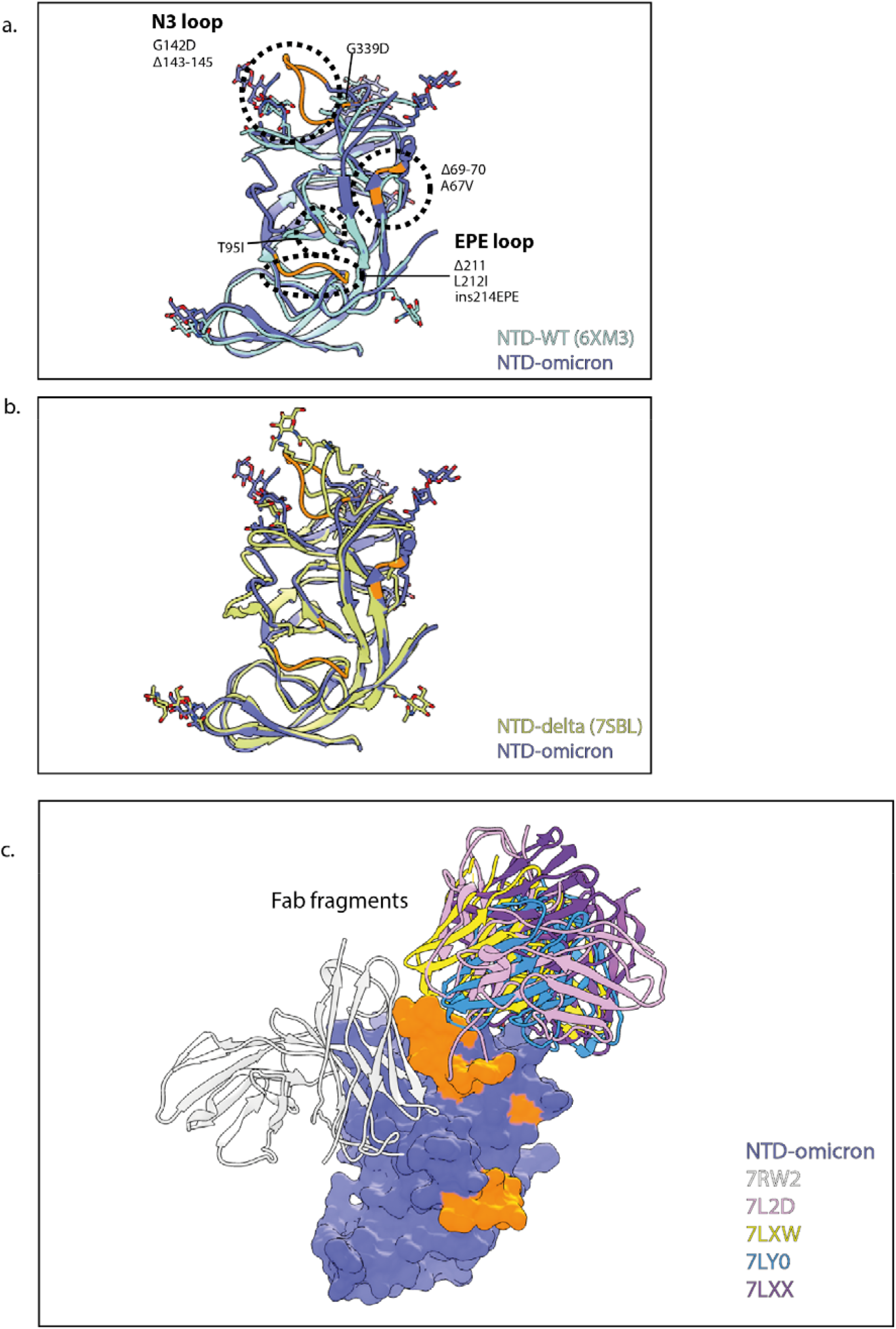
The NTD of the Omicron Spike. Ribbon representations of the Omicron NTD. Colors of models represent the Omicron Spike monomer chain as colored in Fig. 1a. a. Superposition of the Omicron NTD with a wild-type NTD. Orange indicates sites of mutations and loops whose amino acid residues have changed. Circled are the N3 loop containing mutations G142D and deletions 143-145, and the EPE loop containing deletion 211, L212I and insertion at 214 of EPE. b. Superposition of Omicron NTD with a Delta NTD. Coloring is as panel a. c. Structures of NTD neutralizing Fab complexes bound to NTD and superimposed on the Omicron NTD. The Omicron NTD is shown as a surface representation and the Fabs are shown as ribbons. The sites of mutation are indicated and colored as in panel a.

**Figure S5:**
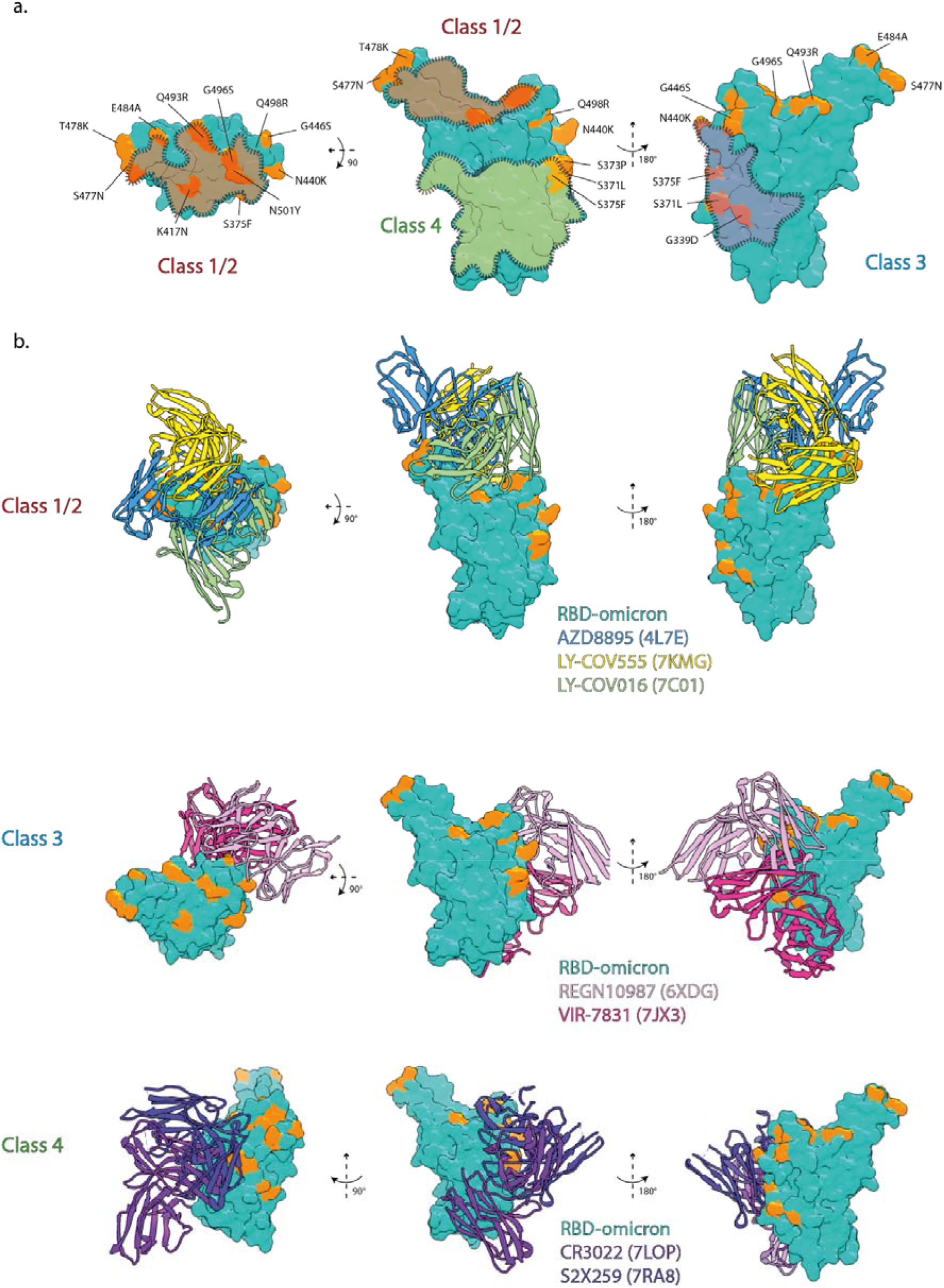
Antibody immune evasion by the Omicron RBD. Ribbon representations of the Omicron RBD. Colors of models represent the Omicron Spike monomer chain as colored in Fig. 1a. a. Surface representation of the Omicron RBD shows in different views with mutations colored in orange. Highlighted with transparent surfaces are the binding surfaces of class 1/2, 3 and 4 antibodies that overlap with Omicron mutations. b. Superpositions of class 1/2, 3 and 4 Fab-RBD structures with the Omicron RBD. Representative Fabs from each class would overlap with the mutated surfaces.

**Figure S6:**
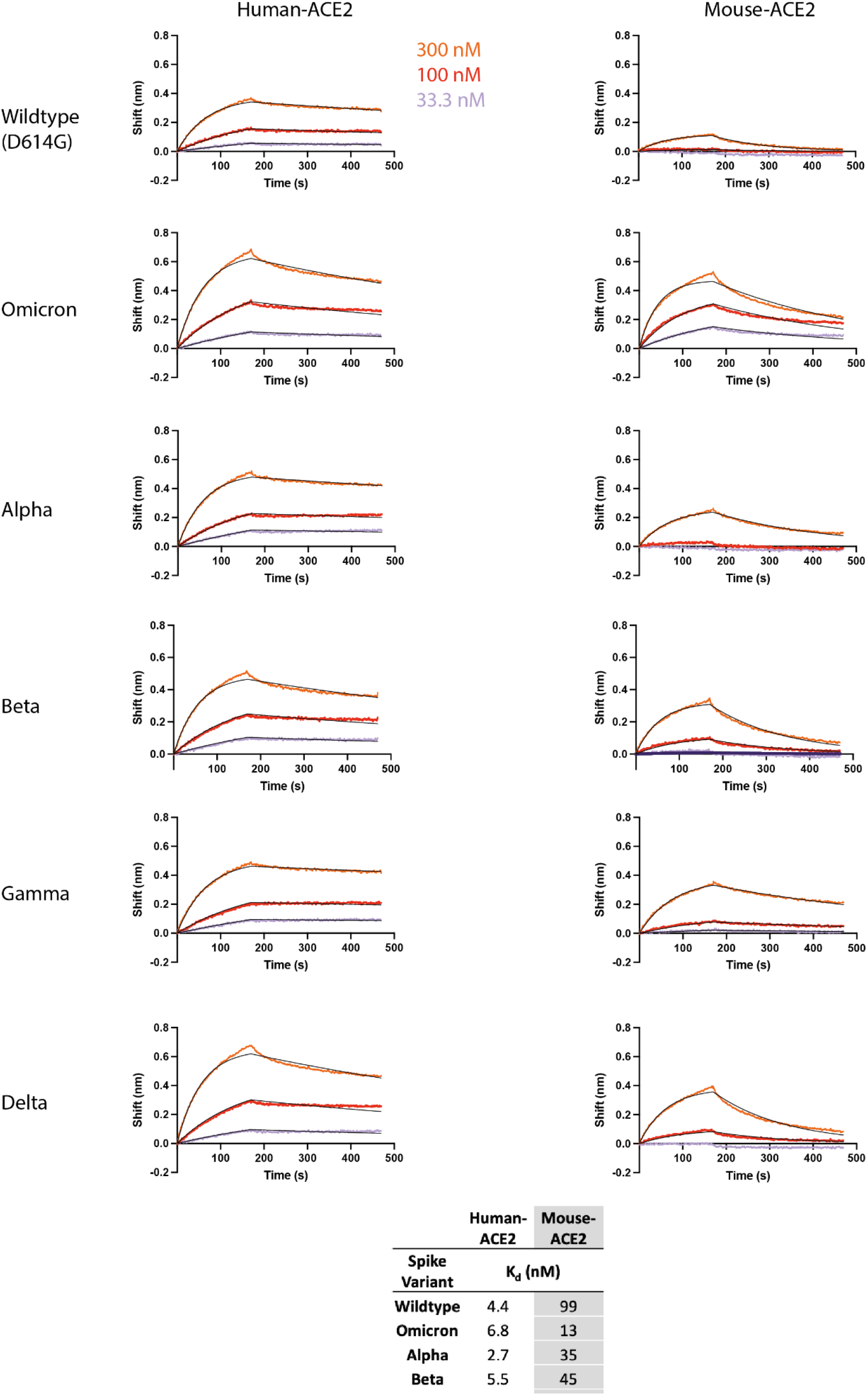
Dimeric human and mouse ACE2 binds to the full-length Omicron SPIKE. BLI curves of indicated Spike binding to captured dimeric human or mouse ACE2. Raw data is colored by concentration as indicated. The black lines represent the fit to the raw data using a 1:1 binding model.

## METHODS

### Protein Production and Purification

RNA isolated from an anonymized leftover sample of a SARS-CoV-2 infected individual was reverse transcribed into cDNA and the Omicron Spike ectodomain was amplified by PCR, sequence verified and cloned into the nCoV-2P-F3CH2S plasmid, replacing the original wildtype Spike.^5^ Mutations stabilizing the Spike protein in the trimeric prefusion state were introduced simultaneously by PCR and In-Fusion cloning. The final construct encodes the Omicron Spike ectodomain, containing a native signal peptide, residues 986 and 987 mutated to proline (2P), a mutated furin cleavage site (residues 682-685 mutated to GSAS), a C-terminal T4 foldon fusion domain to stabilize the trimer complex followed by C-terminal 8x His and 2x Strep tags for affinity purification.^5^ The trimeric Spike protein was transiently expressed in 200 mL of suspension-adapted ExpiCHO cells (Thermo Fisher) in ProCHO5 medium (Lonza) at 5 x10^6^ cells/mL using PEI MAX (Polysciences) for DNA delivery. At 1 h post-transfection, dimethyl sulfoxide (DMSO; AppliChem) was added to 2% (v/v). Following a 2.5-day incubation with agitation at 31 °C and 4.5% CO2, the cell culture medium was harvested and clarified using a 0.22 μm filter. The conditioned medium was loaded onto a 1 mL Streptrap HP column (Cytiva) with a P1 pump, washed with PBS and eluted with 10 mM desthiobiotin in PBS. Eluted protein exchanged into PBS via centrifugal concentrators to 40 uL. Final concentration was 0.35 mg/mL. Final yield was 70 ug/L of culture. The purity of Spike trimers was determined to be >99% pure by SDS-PAGE analysis. (Fig. S1) The protein eluted as a highly-glycosylated trimeric species as determined by SEC-MALS analysis.

### Cryo-electron microscopy

Cryo-EM grids were prepared with a Vitrobot Mark IV (Thermofisher Scientific (TFS)). Quantifoil R1.2/1.3 Au 400 holey carbon grids were glow-discharged for 120 s at 15mA using a PELCO easiGlow device (Ted Pella, Inc.). 3.0 μL of a 0.35 mg/ml protein sample was applied to the glow-discharged grids, and blotted for 6 s under blot force 10 at 100% humidity and 4 °C in the sample chamber, and the blotted grid was plunge-frozen in liquid nitrogen-cooled liquid ethane.

Grids were screened for particle presence and ice quality on a TFS Glacios microscope (200kV), and the best grids were transferred to TFS Titan Krios G4. Cryo-EM data was collected using TFS Titan Krios G4 transmission electron microscope (TEM), equipped with a Cold-FEG on a Falcon IV detector in electron counting mode. Falcon IV gain references were collected just before data collection. Data was collected using TFS EPU v2.12.1 using aberration-free image shift protocol (AFIS), recording 4 micrographs per ice hole.

Movies were recorded at magnification of 96kx, corresponding to the 0.83Å pixel size at the specimen level, with defocus values ranging from −0.8 to −1.8 μm. Exposures were adjusted automatically to 60 e^-^/Å^2^ total dose, resulting in an exposure time of approximately 3 seconds per movie. In total, 8’672 micrographs in EER format were collected.

### Cryo-EM image processing

On-the-fly processing was first performed during data acquisition for evaluating the data quality during screening by using cryoSPARC live v3.3.1.^4^ The obtained ab-initio structures were used for better particle picking for template creation. Motion correction was performed on raw stacks without binning using the cryoSPARC implementation of motion correction.^32^ 470’735 particles were template-based automatically picked and particles were binned by a factor of 4. Two rounds of 2D classification were performed, resulting in a particle set of 144’425 particles. Selected particles resulting from the 2D classification were used for ab-initio reconstruction and hetero-refinement. After hetero-refinement, 76’450 particles contributed to an initial 3D reconstruction of 3.02 Å resolution (FSC 0.143) with C1 symmetry, while application of C3 symmetry resulted in a 3D map at 2.65 Å resolution (FSC 0.143). The same number of particles were then subjected to local 3D refinement. A soft mask volume against the RBD-NTD area was generated manually in UCSF Chimera and cryoSPARC.^33^ The RBD-NTD region was subtracted from the global trimeric spike and further focused refinement was performed subsequently, which resulted in a 3D map at 3.88 Å resolution (FSC 0.143).

### Cryo-electron microscopy model building

A model of a Spike trimer (PDB ID 6MXL) or AlphaFold2 (ColabFold implementation) models of the Spike’s NTD and RBD were fit into the cryo-EM maps with UCSF Chimera.^15^ These docked models were extended and rebuilt manually with refinement, using Coot and Phenix.^34,35^ Figures were prepared in UCSF Chimera and UCSF ChimeraX.^33^

### Biolayer Interferometry (BLI)

Running buffer was 150 mM NaCl, 10 mM HEPES 7.5 and all experiments were performed on a Gator BLI system. Dimeric mFc ACE2 was diluted to 10 ug/mL and captured with MFc tips (GatorBio). Loaded tips were dipped into a 3-fold serial dilution series (300 nM, 100 nM, 33.3 nM) of analyte SPIKE. Curves were processed using the Gator software with a 1:1 fit after background subtraction.

## DATA AVAILABILITY

The reconstructed maps of the full Omicron Spike are available from the EMDB database, C1 symmetry, EMDB-14086. The atomic model for the full-length Omicron Spike is available from the PDB database, PDB-7QO7. The local focused-refinement map of the NTD and RBD regions are available from the EMDB database, EMDB-14087. The atomic model for the NTD and the RBD in the locally refined map is available from the PDB database, PDB-7QO9. The raw cryo-EM images will be deposited in the EMPIAR database.

## CONTRIBUTIONS

P.T. and C.R. isolated and cloned the Omicron Spike gene and prepared DNA for protein expression. K.L. purified the Omicron Spike. B.B., A.M., and S.N., froze and screened cryo-EM grids, collected data, and performed on-the-fly processing. D.N. processed cryo-EM data and determined the final model. D.N. and K.L. analyzed data and prepared figures. K.L., D.N., F.P., H.S., and D.T. wrote the manuscript. F.P., A.M., H.S., and D.T. supervised the project.

## ACKNOWLEDGEMENTS

Cryo-EM data collection and initial image processing was performed at the Dubochet Center for Imaging, a common initiative of the EPFL, UNIL and UNIGE. We thank Rosa Schier, Laurence Durrer, Soraya Quinche and Michael François of the Protein Production and Structure Core Facility of EPFL for helping in the production of Spike Omicron. We thank Isabella Eckerle, Meriem Bekliz and the Virology laboratory of the HUG for the RNA sample collection. This work was supported by the Swiss National Science Foundation, grant NCCR TransCure.

## Notes

### Competing Interest Statement

The authors have declared no competing interest.

### Summary of Updates

Revision 2 - submitted in error. Was for another manuscript. Back to Revision 1 version.

